# Inhibition of JEV infection using β-Catenin specific inhibitor, iCRT-14

**DOI:** 10.64898/2026.08.29.747967

**Authors:** Ankita Datey, Soumyajit Ghosh, Sanchari Chatterjee, Bijita Bhowmick, Archana Ghatak, Bharat Bhusan Subudhi, Soma Chattopadhyay

## Abstract

The lack of effective anti-JEV therapy possesses significant challenge to control JEV. β-catenin, a key mediator of Wnt signaling pathway regulates different viral replication and host immune responses. However, its role in JEV infection remains to be elucidated. Thus, the current study focused on evaluating iCRT-14, a specific β-catenin inhibitor, against JEV. Treatment with iCRT-14 following JEV infection resulted efficient reduction in viral progeny release, viral RNA and protein levels in Huh7 and HEK293T cells. Further, active and total β-catenin, Cyclin D-1 and GSK3-β, the other key pathway players were also modulated in infected and inhibitor treated cells. Moreover, iCRT-14 showed an IC□ □ of 4.56 µM in Huh7 cell and maximal inhibition at the early stages of the JEV life cycle. Interestingly, the overexpression of β-catenin in both the cells and siRNA-mediated β-catenin knockdown (in Huh7 cells) significantly abrogated JEV replication, as evidenced by decreased viral titers, viral protein expression, and viral as well as total RNA levels. Moreover, the reduction in extracellular (84%) and intracellular (60%) viral titers following iCRT-14 treatment highlights its role in impairing JEV infection. Further, *in silico* molecular docking and co-immunoprecipitation studies demonstrated interactions between β-catenin and the JEV NS5 and E proteins. Collectively, these findings suggest that optimum level of β-catenin is required for efficient JEV infection, highlighting its potential as a target for designing host-directed control strategies to regulate viral infection.

## 1. Introduction

Japanese Encephalitis Virus (JEV), a flavivirus transmitted by *Culex* mosquitoes, persists to be a leading cause of acute viral encephalitis in many regions of Asia and the Western Pacific. With the ability to cross the blood-brain barrier (BBB), JEV primarily impact children and immunocompromised individuals. This leads to a spectrum of complications ranging from mild febrile illness to severe neurological complications (Zhu et al., 2023). The 11 kb long single-stranded positive-sense RNA genome of JEV encodes for three structural (Capsid, Envelope and Membrane) and seven non-structural (NS) proteins vital for viral replication and immune modulation (Guo et al., 2020). Although only a small proportion of JEV infections become symptomatic, the virus remains a major public health concern in endemic regions, causing an estimated 100,000 clinical cases annually, with mortality rates reaching up to 30% among encephalitis patients, and prolonged neurological sequelae affecting nearly half of the survivors (https://www.who.int/news-room/fact-sheets/detail/japanese-encephalitis).

Despite the availability of effective vaccines against JEV, their implementation remains suboptimal in the low- and middle-income countries due to geographical, economic, and regulatory barriers (Pilihanto et al., 2025). Also, the absence of clinically approved drugs or antiviral therapies create challenges in complete disease control (Zhu et al., 2023). This emphasizes the pressing need to explore alternative host factor mediated antiviral approaches. One such potentially emerged essential host protein is β-catenin, which functions as a key mediator of Wnt signaling pathway. This is involved in gene transcription, proliferation of cells and immune regulation (Chatterjee et al., 2023). Several evidences have implicated β-catenin as a modulator of viral replication and host immune responses in different viral infections including Chikungunya (CHIKV), Influenza A (IAV), Dengue and Zika viruses (Chatterjee et al., 2023; Hillesheim et al., 2014; Jimenez et al., 2021; Smith et al., 2017). The role of β-catenin as junction proteins have been reported earlier during JEV infection (Agrawal et al., 2013), however, much remain to be explored.

iCRT-14, initially explored for anticancer properties, is a potent inhibitor of β-catenin-responsive transcription (CRT). This drug attenuates β-catenin-mediated gene expression by disrupting its interaction with T-cell factor/lymphoid enhancer-binding factor (TCF/LEF) which are transcriptional regulatory proteins (More et al., 2018). Considering the emerging significance of β-catenin in different viral infections, the current study aims to investigate the effect of iCRT-14 in JEV infection which might help in designing novel therapeutic approaches.

## 2. Materials and methods

### 2.1. Cells and Virus

Huh7 (Human Hepatocarcinoma; NCCS, Pune), PS (Porcine stable kidney) and HEK293T (Human embryonic kidney; NCCS, Pune) cells were maintained in Dulbecco’s Modified Eagle Medium (DMEM; Himedia, India) supplemented with 10% fetal bovine serum (FBS; Gibco, USA), penicillin-streptomycin (Himedia), and gentamycin (Gibco, USA). The GP78 strain of JEV (accession No. **AF075723**) as well as PS cells were kindly provided by Dr. Anirban Basu, National Brain Research Centre (NBRC), Gurgaon, India.

### 2.2. Cytotoxicity assay

The cytotoxicity of iCRT-14 (SML0203; Sigma Aldrich, USA) was evaluated using the EZcount MTT cell assay kit (CCK003; Himedia, India) as described before (Datey et al., 2025). Huh7 and HEK293T cells were treated with varying concentrations of the inhibitor for 24□hours (h). Next, MTT solution was added to the cells and incubated for 2 h, followed by the addition of solubilization solution. The absorbance was measured at 570□nm using a multimode plate reader (PerkinElmer, MA) and cell viability was estimated accordingly.

### 2.3. Virus infection

The JEV [MOI (multiplicity of infection): 0.1 or 0.01] infection was performed in both the Huh7 and HEK293T cells in serum-free media (SFM) with shaking at every 10 to 15□minutes (min) intervals for 90 min, as stated earlier (Datey et al., 2025).

### 2.4. Plaque Assay

Plaque Assay was performed as described previously (Datey et al., 2025) to determine viral titers [as plaque-forming units per milliliter (PFU/mL)]. In brief, culture supernatants from different treatment conditions were serially diluted and added to PS cell monolayers, followed by incubation for 1.5 h. Next, cells were washed with 1X phosphate-buffered saline (PBS) and overlaid with methylcellulose (Sigma, USA). After 6-7 days, cells were fixed with 8% formaldehyde, stained with 0.1% crystal violet (Sigma) and the resulting plaques were counted. Further, to assess whether the inhibitor interferes in viral replication, intracellular and extracellular viral titres were measured as described before (Lee and Nishizawa, 2024).

### 2.5. Western blot

The western blot analysis was carried out as reported earlier (Chatterjee et al., 2023). Briefly, the cells were lysed with Radio Immuno Precipitation Assay (RIPA) buffer and the protein lysates were separated by 10% SDS-polyacrylamide gel, and transferred onto PVDF membranes (Polyvinylidene Fluoride). The membranes were then probed with primary antibodies against total/active β-catenin [ab16051, Abcam; 8814S, Cell Signaling Technology (CST), USA], GSK-3β (Glycogen synthase kinase-3beta; 12456S, CST), JEV-NS3, NS5 and Envelope (GTX125868, GTX131359 and GTX125867 respectively, GeneTex, USA). Glyceraldehyde 3-phosphate dehydrogenase (GAPDH; AC002, ABclonal, USA) was used as loading control. The blots were developed using the Immobilon Western Chemiluminescent HRP substrate (Millipore, USA) and band intensities were quantified using the ImageJ software.

### 2.6. qRT-PCR

Total RNA and viral RNA were extracted following the earlier method (Datey et al., 2025) and quantitative reverse transcription PCR (qRT-PCR) was performed using Luna Universal qPCR Master Mix (M3003L, New England Biolabs, USA) with JEV-Envelope, GAPDH and Cyclin D-1–specific primers (Table S1). The relative fold change and viral RNA copy number/mL were calculated as described before (Datey et al., 2025).

### 2.7. Inhibitor treatment at different phases of viral infection

The pre, during and post-treatment experiment with iCRT-14 was carried out as mentioned before with little modifications (Mahapatra et al., 2025). In short, in pre-treatment condition the cells were incubated with the inhibitor for 3 h and then infection was carried out. In during-treatment condition, the inhibitor was added to the cells during infection, whereas in the post condition the inhibitor was added to cells after infection. For the combined (pre-during-post) treatment, the drug was present before, during, and after infection. The supernatants from the respective treatment conditions were collected at 24 hpi and subjected to plaque assay.

### 2.8. Time-of-addition experiment

In the time-of-addition experiment, the inhibitor was added to JEV infected cells separately at 0, 4, 8, 12, 16, 20, and 24 hpi (hours post infection). At 26 hpi, the supernatants were harvested to estimate the viral titer as described previously (Chatterjee et al., 2023).

### 2.9. Determination of half-maximal inhibitory concentration (IC_50_)

For determining the IC_50_ of iCRT-14, JEV infected cells were treated with different concentrations (2, 4, 8 and 10µM) of the inhibitor. The supernatants were collected at 24 hpi and plaque assay was performed. The nonlinear regression analysis was performed to fit the data and the concentration of iCRT-14 required to inhibit 50% of the viral burden was calculated as mentioned previously (Mahapatra et al., 2025).

### 2.10. siRNA or plasmid transfection

Huh7 cells were transfected with either 90 picomolar (pM) of β-catenin siRNA (Table S1) or with scrambled siRNA using Lipofectamine 3000 reagent (L3000008; Invitrogen, USA) as per manufacturer’s protocol. At 48 h post-transfection (hpt), cells were infected with JEV (MOI of 0.1) and supernatants and cells were collected at 24 hpi for plaque assay and Western blotting as reported earlier (Datey et al., 2025). For overexpression studies, 1µg of β-catenin plasmid (Addgene, USA) was transfected similarly into both Huh7 and HEK293T cells, where empty vector (pcDNA3) was used as a negative control. Subsequently, JEV infection (0.1 MOI) was carried out in the transfected cells as described previously (Chatterjee et al., 2023). Culture supernatants and cells were collected at 24 hpi and subjected for downstream experiments.

### 2.11. Co-Immunoprecipitation (CO-IP)

For co-immunoprecipitation, mock and JEV-infected Huh7 cells were harvested at 24 hpi and lysates were processed using the Dynabeads Protein A Immunoprecipitation Kit (10006D; Invitrogen, USA) following the earlier protocol (Chatterjee et al., 2023).

### 2.12. Molecular docking

The protein-protein interaction studies were performed using the Cluspro 2.0 web server as described earlier (Nayak et al., 2019). In short, high resolution protein structures (PDB ID: 2Z83 for NS3 helicase, 4K6M for NS5MTase, 4HDH for NS5RdRp, 5036 for NS1, 5MV1 for E and 8Z61 for human β-catenin) were obtained from the PDB database and processed using the Discovery studio program. The protein-protein docking was carried out using the ClusPro2.0 webserver. The most stable complex (balanced mode) was retrieved and processed in the PyMol software. This complex was further evaluated using the HyPPI program of the Protein plus webserver to predict the nature of interactions. Also, complex binding properties were analysed using the Prodigy Webserver and the interaction diagram created using the Maestro program.

### 2.13. Statistical analysis

Statistical significance was determined by the one-way ANOVA (Brown-Forsythe) or unpaired Student’s t-test using the GraphPad Prism 8.0.1 software. Data are presented as mean ± standard deviation (SD) with n=3. P values were reported as *≤0.033, **≤0.002, ***≤0.0002, ****≤0.0001.

## 3. Results

### 3.1. Treatment with iCRT-14 diminishes JEV infection significantly *in vitro*

In order to assess the effect of iCRT-14 during JEV infection, MTT assay was performed to determine its cytotoxicity in the Huh7 cells. The result demonstrated that iCRT-14 was non-cytotoxic at concentrations up to 75□µM (Fig 1A). Next, the anti-JEV property of the inhibitor was evaluated in JEV infected (0.1 MOI) Huh7 cells. Post infection, treatment with 10□µM of iCRT-14 resulted >96% reduction in viral plaques compared to the infection control (Fig 1B). Further, there was a significant decrease (68%) in total RNA level of JEV-E gene in the inhibitor treated condition as compared to infection only (Fig 1C). Notably 30%, 48% and 30%, 55% reduction was observed in total and active β-catenin levels in infected and iCRT-14 treated samples, respectively, compared to mock (Fig 1D-F). Moreover, around 60% reduction was noticed in JEV-NS3 protein level in treated condition compared to the infection control (Fig 1D and G). The inhibition was more robust when the cells were infected at a lower MOI of 0.01 as JEV titer and JEV-NS3 levels were reduced by ∼99% and 75%, respectively in presence of iCRT-14 (Fig S1A-C). In addition, the total β-catenin protein was reduced by 50% after infection and 71% upon treatment, in comparison with mock (Fig S1B and D).

**Figure 1:**
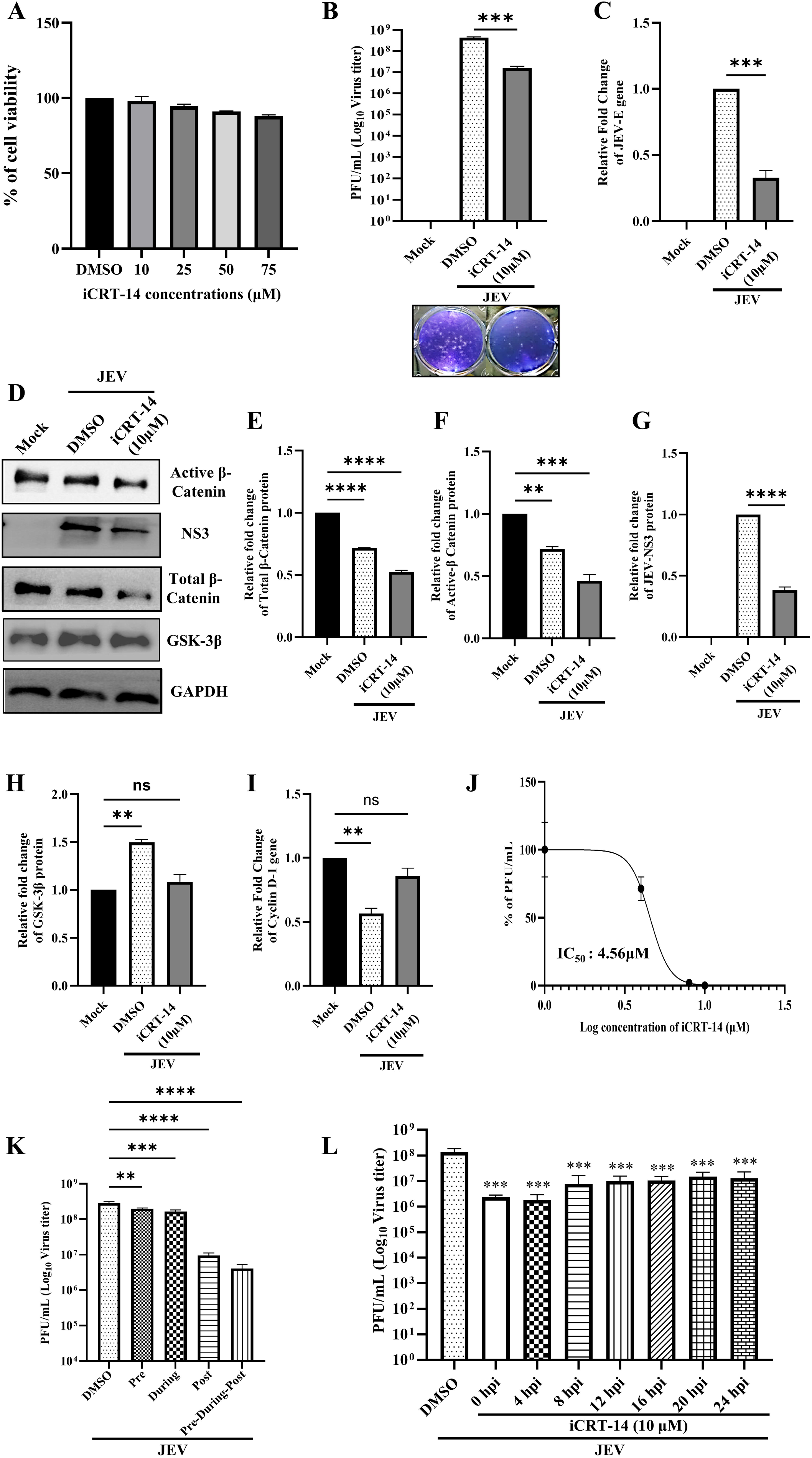
Inhibition of JEV infection by iCRT-14 *in vitro*. Huh 7 cells were treated with different concentrations (10, 25, 50 and 75µM) of iCRT-14 for 24 h and by MTT assay the cytotoxicity of the inhibitor was assessed. **(A)** Graph showing viability of cells. **(B)** Huh 7 cells were infected with JEV (MOI of 0.1) and iCRT-14 (10 µM) was added post infection. The supernatants were collected at 24 hpi and subjected to plaque assay to determine the viral titers in infected and inhibitor treated samples. Bar graph representing the log_10_ of viral titer (PFU/mL). **(C)** Total RNA was isolated from the mock, JEV infected and treated cells. The JEV-E gene was amplified by qRT-PCR. Bar diagram demonstrating fold changes of JEV-E gene. **(D)** Images representing expression levels of active β-catenin, JEV-NS3, total β-catenin and GSK-3β proteins, estimated through Western blotting in the mock, infected and treated samples respectively. GAPDH was served as a loading control. **(E-H)** Bar diagrams depicting the relative fold changes of total and active β-catenin, JEV-NS3 and GSK-3β proteins. **(I)** Bar graph showing changes in expression of Cyclin D-1 gene in mock, infected and treated samples. **(J)** The IC_50_ value of iCRT-14 was determined against JEV in Huh7 cells and plotted as logarithmic values of various doses of iCRT-14 on the X-axis and % of viral titer (PFU/mL) on the Y-axis. **(K)** Huh7 cells were treated with the inhibitor at pre (3 h prior to infection), during (90 minutes during infection) and post infection (for 24 h) conditions. The inhibitor was present before, during, and after infection for the combined treatment (pre-during-post) condition. Bar diagram demonstrating the log_10_ of JEV titer of the supernatants, collected at 24 hpi. **(L)** Huh7 cells were infected with JEV (0.1 MOI) and iCRT-14 was added to the cells at every 4 h intervals from 0 to 24 hpi. Bar diagram representing log_10_ of viral titers of all the supernatant samples, collected at 26 hpi. Data of three independent experiments are shown as mean ± SD with P values: *<0.03, **≤0.002, ***≤0.0002, ****<0.0001 were considered statistically significant.

GSK-3β, a negative regulator of β-catenin, facilitates its ubiquitination and degradation by participating in the destruction complex. Further, β-catenin regulates key downstream targets including Cyclin D-1, c-MYC and IFN-β in which Cyclin D-1 specifically controls progression through the G1 phase of the cell cycle (Du et al., 2020). Therefore, in the present study the expression levels of Cyclin D-1 and GSK-3β proteins were evaluated. The GSK-3β expression was upregulated by 51% in infected samples as compared to mock, whereas downregulated by 45% in iCRT-14 treated samples, relative to the infection condition (Fig 1H). Also, 44% reduction was observed in the expression of Cyclin D-1 gene of infected samples as that of mock, whereas it showed significant upregulation in iCRT-14 treated samples (Fig 1I). Similar observations were found when HEK293T cells were infected with JEV (0.1 MOI) and treated subsequently with a non-toxic dose of the inhibitor (10□µM; Fig S2A). The total beta catenin was found to be downregulated in infected (27%) and iCRT-14 (45%) treated cells significantly (Fig S2B and C). Also, a remarkable reduction (57%) in JEV-NS3 was observed in the iCRT-14 treated cells as relative to that of infection control (Fig S2B and D). Moreover, the inhibitor-treated samples exhibited a one-log reduction in viral RNA levels (copy number/mL) compared to the infection control (Fig S2E). The IC_50_ value of iCRT-14 against JEV in Huh7 cells was found to be 4.56 µM (Fig 1J). Further, to evaluate whether the inhibitor interferes in the viral entry/attachment or replication processes, pre-, during-, and post-treatment experiments were performed in both Huh7 and HEK293T cells as mentioned above. In case of Huh7 cells, pre-treatment with iCRT-14 resulted in 31% reduction in viral titer. Whereas, 43% reduction was observed in the during treatment condition and post-treatment led to >96% reduction in viral titer compared to the infection control. Interestingly, the combined (pre-during-post) treatment revealed maximum inhibition (∼99%) in viral progeny release (Fig 1K). In the HEK293T cells, pre, during, and post-treatment conditions demonstrated 33%, 39% and 89% reduction in JEV RNA copies, respectively (Fig S2F). However, highest reduction in viral RNA (96%) was also observed upon the combined treatment (Fig S2F).

As more reduction was observed in post iCRT-14 treated condition (Fig 1K), a ‘time of addition’ experiment was performed to determine at which stage of the viral life cycle β-catenin is involved. It was observed that iCRT-14 was able to inhibit viral progeny release at all phases of post infection condition, however the effect was more pronounced during early phase (96-99% reduction) (Fig 1L).

Thus, collectively these findings suggest that iCRT-14, a β-catenin inhibitor, can abrogate JEV infection significantly.

### 3.2. JEV requires optimal β-catenin level for efficient infection

In order to understand the role of β-catenin during JEV infection, modulation of β-catenin was performed in Huh7 cells, either by overexpression or siRNA-based knockdown experiments. As observed in Fig 2A and B, the overexpression of β-catenin after transfecting the plasmid resulted in significant increase in β-catenin level (43%) to that of empty vector control. It led to 50% reduction in the JEV-NS3 protein level (Fig 2A and C). The viral titer was also decreased significantly (80%) upon overexpression, as compared to the empty vector control (Fig 2D). Further, β-catenin expression was silenced using siRNA (90pM) and the knockdown efficiency was confirmed by Western blot analysis. It was observed that total β-catenin level was reduced by 40% as compared to the scramble control (Fig 2E and F). Subsequently, the siRNA-transfected cells were infected with JEV and protein expressions were assessed at 24 hpi by Western blotting. The JEV-NS3 as well as total β-catenin expression levels were decreased by 42% and 53% respectively, compared to scramble control (Fig 2G-I). Similarly, 95% reduction was observed in the viral titer of siRNA knockdown samples compared to that of scramble only (Fig 2J). Further, the β-catenin overexpression data in HEK293T cells revealed 30% depletion of the JEV-NS3 protein level in beta catenin overexpressed (28%) cells as compared to the empty vector control (Fig S2G-I). The viral RNA was also significantly reduced (one-log reduction) in the beta catenin overexpressed HEK293T cells, when compared with the empty vector control (Fig S2J).

**Figure 2:**
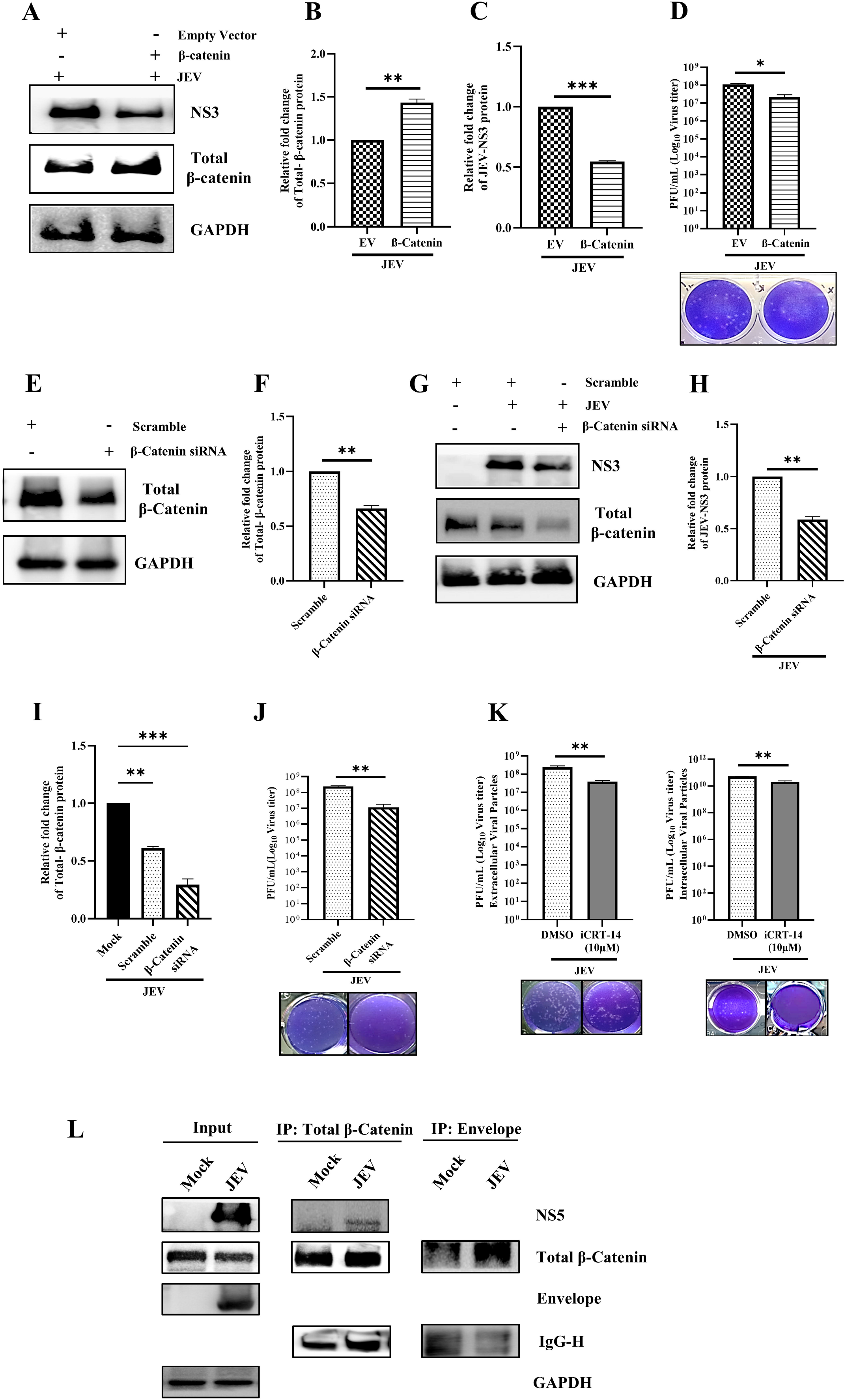
iCRT-14 might act by hindering JEV replication machinery. Huh 7 cells were transfected with β-catenin plasmid for its overexpression and subsequently infected with JEV (MOI 0.1) at 24 h post transfection (hpt). The cells and culture supernatants were harvested at 24 hpi for downstream experiments. **A)** Western blot images showing the levels of total β-catenin and JEV-NS3 proteins. GAPDH served as loading control. **(B and C)** Bar diagrams showcasing relative fold changes of total β-catenin and JEV-NS3 protein levels. **(D)** Bar graph depicting the log_10_ of viral titer (PFU/mL) in the culture supernatants (empty vector + JEV and overexpression of β-catenin + JEV). **(E)** Huh 7 cells were transfected with scrambled siRNA or β-catenin siRNA (90pM) for 48 h. Western blot image showing the total β-catenin protein levels after siRNA transfection. **(F)** Graph representing relative fold change of total-β-catenin upon siRNA-mediated knockdown. **(G)** Western blot images depicting total β-catenin and NS3 proteins after transfection and JEV infection. **(H and I)** Bar graphs showing relative fold changes of JEV NS3 and total β-catenin proteins. **(J)** Bar diagram representing viral titer in log_10_ scale in the culture supernatants of scramble control and siRNA (90pM) treated along with JEV infected samples. **(K)** Bar graph showing extracellular and intracellular viral particles as log_10_ PFU/mL in infected and iCRT-14 treated samples. **(L)** Huh7 cells were infected with JEV for 24 h. The cell lysates were co-immunoprecipitated with either total β-catenin or JEV-E antibodies. The western blot images showing the levels of JEV-NS5, total β-catenin, JEV-E and GAPDH in the whole cell lysate (left panel). Middle panel represents the interaction of total β-catenin with JEV-NS5 protein, whereas the interaction of JEV-E and total β-catenin is shown in the right panel. Data are expressed as mean ± SD from three independent experiments. Statistical significance was defined as P values *≤0.03, **≤0.002, ***≤0.0002.

Additionally, to understand whether iCRT-14 impairs JEV replication, extra and intracellular viral titers were evaluated post inhibitor treatment. It was observed that both extracellular and intracellular viral titers showed 84% and 60% decrease respectively in treated conditions compared to infection control (Fig 2K). Furthermore, to understand the potential interactions between β-catenin and JEV proteins, molecular docking analyses were performed using the viral E, NS1, NS3 and NS5 proteins. The data revealed that JEV-NS5 (RdRp) and E proteins exhibited greater likelihood of complex formation with β-catenin compared to the other viral proteins (Table S2). Next, to validate the docking result, Huh7 cells were infected with JEV and subjected to co-immunoprecipitation. It was observed that the JEV NS5 and E proteins were interacting with the total β-catenin protein (Fig 2L).

Taken together, it can be stated that an optimal level of β-catenin might be required for efficient JEV replication during its initial phase of life cycle. This might be through its interaction with different viral proteins.

## 4. Discussion

The unavailability of anti-JEV therapeutics continues to pose a significant challenge to control JEV cases. Although several vaccines are available, factors such as high cost, the requirement for multiple doses, the emergence of new viral genotypes, and poorly coordinated vector and disease control programs continue to limit the prevention and control of JEV infection. Hence, the current study aims to understand the impact of iCRT-14, a specific inhibitor of β-catenin on JEV pathogenesis.

Here, Huh7 and HEK293T cells were infected with JEV and iCRT-14 was administered post infection which resulted in significant decrease in viral titers, total RNA or viral RNA and the JEV-NS3 protein levels. Further, modulation in the expression levels of the β-catenin pathway proteins such as, active and total β-catenin, Cyclin D-1 and GSK3-β were observed in infected and inhibitor treated samples as compared to mock. The IC_50_ value of iCRT-14 was found to be 4.56 µM. The maximum inhibition of viral progeny release was noticed in the post-treatment condition and the time of addition experiment revealed that iCRT-14 was abrogating the viral progeny release more effectively in the early stages of JEV life cycle.

Additionally, both overexpression and siRNA mediated knockdown of β-catenin resulted in significant decrease in JEV particle formation and NS3 levels. Moreover, extracellular and intracellular viral titers were reduced remarkably (84% and 60% respectively) after iCRT-14 treatment. Further, *in silico* analysis demonstrated that β-catenin can interact with the JEV-NS5 and E proteins which were validated experimentally by co-immunoprecipitation.

Previously, similar observations have been reported where viral load was decreased following iCRT-14 treatment in other viral infections such as Chikungunya (CHIKV), influenza and SARS-CoV-2 viruses (Chatterjee et al., 2023; More et al., 2018; Xu et al., 2024). While, in Bovine herpes virus type 1 (BoHV-1); iCRT-14 administration enhanced infection (Ding et al., 2022). Earlier reports further suggested that β-catenin expression was reduced during CHIKV, ZIKV and BoHV-1 infections (Chatterjee et al., 2023; Ding et al., 2022; Jimenez et al., 2021), which are similar to the current observation. In contrary, SARS CoV-2 and Pseudorabies virus (PRV) infection enhanced β-catenin expression for efficient viral proliferation (Wang et al., 2022; Xu et al., 2024), indicating different roles of the Wnt/β-catenin signaling pathway in various viral infections. In addition to β-catenin, previous studies have demonstrated that JEV-induced downregulation of Cyclin D1 results in cell-cycle arrest (Kim et al., 2015), corroborating the observations of the present study. The increased Cyclin D-1 levels following iCRT-14 treatment may indicate partial restoration of this arrest, potentially through a β-catenin–independent mechanism. Changes in GSK-3β levels observed during JEV infection and iCRT-14 treatment also align with reports from CHIKV and Bovine parainfluenza virus type 3 (BPIV3) infections (Chatterjee et al., 2023; Du et al., 2020). Thus, β-catenin downregulation in JEV infection might occur through a GSK-3β mediated pathway (Du et al., 2020) that requires further investigation.

In case of JEV, iCRT-14 was effective in post and to some extent in both pre and during treatment conditions, indicating its interference in viral replication and/or entry processes. Previous reports also demonstrated inhibition of CHIKV and PRV infections following iCRT-14 treatment (Chatterjee et al., 2023; Wang et al., 2022). Reduction in intracellular and extracellular viruses further supported the role of iCRT-14 in impairing replication (Kasabe et al., 2023). Overall; these data suggest that β-catenin contributes to multiple stages of the JEV life cycle that needs further investigation.

In the present study, both overexpression and knockdown approaches indicate that an optimal level of β-catenin is essential for efficient JEV replication. Importantly, this observation is supported by experiments performed in two different human cell lines (Huh7 and HEK293T). Furthermore, these results provide insight into the potential for developing host-targeted therapeutic approaches. Specifically, targeting β-catenin signaling could have differential effects depending on the basal expression levels in distinct cell types, which needs detailed studies in future.

Earlier it was documented that β-catenin can interact with viral proteins like, nsP2, Nef, UL8, viral nucleoprotein (N) and phosphoprotein (P) of CHIKV, HIV, HCMV, HPIV-3 respectively (Bose and Banerjee, 2004; Chatterjee et al., 2023; Dirck et al., 2023; Weiser et al., 2013). In the present investigation, JEV NS5 and E proteins were also found to interact with β-catenin; however detailed mechanistic insights are yet to be explored.

This study lacks *in vivo* validation which would be important for assessing the preclinical efficacy of iCRT-14 against JEV. Nonetheless, further investigation is needed to clarify the precise role of β-catenin during JEV infection. In conclusion, β-catenin is found to be essential for efficient JEV infection, highlighting its potential as a target for designing host-directed control strategies to regulate viral infection.

## CRediT authorship contribution statement

**Ankita Datey:** Conceptualization, Data curation, Investigation, Methodology, Validation, Writing – original draft; **Soumyajit Ghosh**: Data curation, Formal analysis, Investigation, Methodology, Validation, Writing – review and editing; **Sanchari Chatterjee:** Data curation, Investigation, Validation; **Bijita Bhowmick**: Data curation, Investigation, Methodology, Writing – original draft; **Archana Ghatak:** Resources; **Bharat Bhusan Subudhi:** Formal analysis, Investigation, Software; **Soma Chattopadhyay:** Conceptualization, Funding acquisition, Project administration, Resources, Supervision, Visualization, Writing – review and editing.

## Data availability

The data that support the findings of this study are available in the manuscript itself.

## Acknowledgement

We are thankful to Dr. Anirban Basu for providing the JEV (GP78) virus strain and PS cell line. We would also like to thank Ms. Supriya Suman Keshry and Ms. Anisha Bhattacharya for their assistance during the experiments.

