## Supplemental Data for "Inhibition of JEV infection using β-Catenin specific inhibitor, iCRT-14"

e) Drug Development and Analysis Lab, School of Pharmaceutical Sciences, Siksha 'O' Anusandhan Deemed to be University, Bhubaneswar, Odisha, India.

### Equal contributors

*** Corresponding author**

Address of corresponding author:

Dr. Soma Chattopadhyay
Infectious Disease Biology
BRIC-Institute of Life Sciences
Nalco Square, Bhubaneswar, Odisha, India- 751023
Phone No: +91-674-2304235;


**Keywords**: Japanese encephalitis virus, β-catenin, iCRT-14, GSK3-β

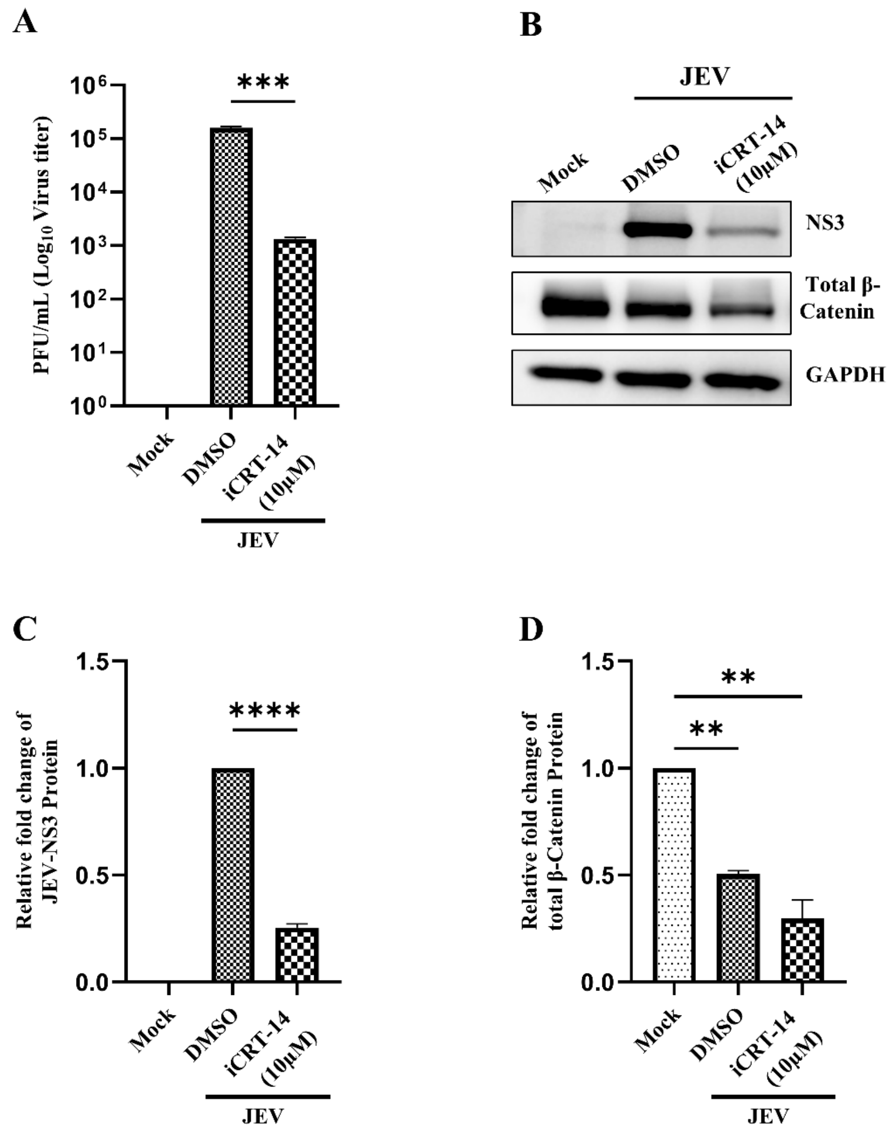


**Figure S1: iCRT-14 restricts JEV infection in Huh7 cells.** Huh 7 cells were infected with JEV (0.01 MOI) and the inhibitor (10 µM) was added post infection. The culture supernatants and cells were harvested at 24 hpi for Plaque assay and Western blotting. (A) Bar diagram representing the $\log_{10}$ of JEV titer (PFU/mL). (B) Westen blot images showing the expression of JEV-NS3 and total β-catenin protein levels in mock, infected and treated samples. (C and D) Bar diagrams indicating the relative fold changes of JEV-NS3 and total β-catenin proteins. Data are presented as mean ±SD (n=3). Differences were considered statistically significant as P values: **≤0.002, ***≤0.0002 and ****<0.0001.

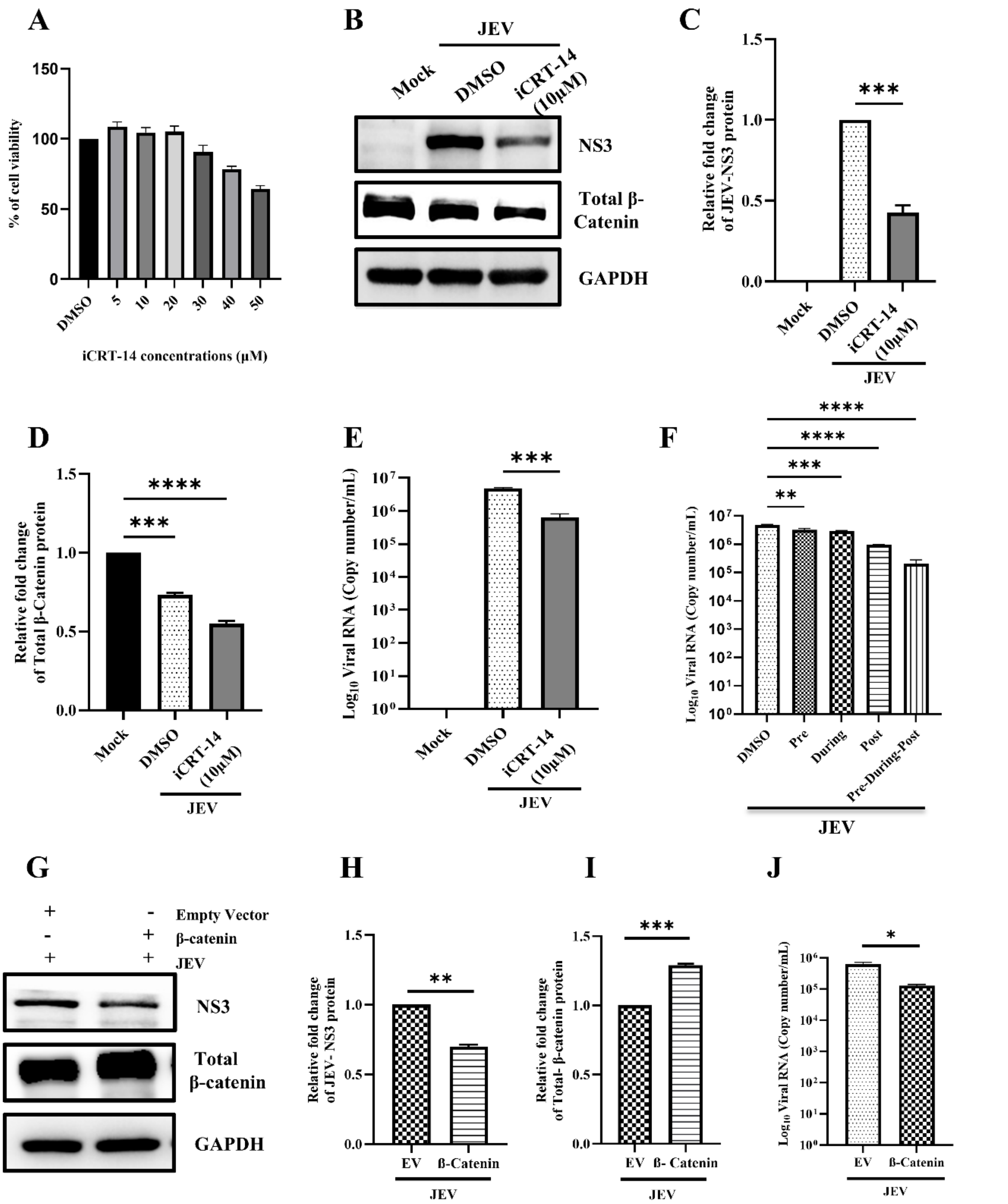


**Figure S2: iCRT-14 abrogates JEV infection in HEK293T cells:** HEK293T cells were treated with different concentrations of iCRT-14 for 24 h and MTT assay was carried out. (A) Bar graph representing percentage of viable cells. (B) HEK293T cells were infected with JEV (MOI 0.1) and treated with iCRT-14 (10 µM) post infection. The cells were harvested at 24hpi and subjected for the Western blotting. Images representing the expression levels of JEV-NS3 and total β-catenin proteins in mock, infected and treated samples. (C, D) Bar Graphs representing the relative fold changes of the JEV-NS3 and total β-catenin proteins. (E) The culture supernatants were also harvested at 24 hpi and subjected to qRT-PCR analysis. Graph representing viral RNA (copy no./mL) in presence and absence of iCRT-14. (F) HEK293T cells were treated with the inhibitor separately at pre (3 h prior to infection), during (90 minutes during infection) and post infection (for 24 h) conditions. The inhibitor was present before, during, and after infection for the combined treatment (pre-during-post) condition. Culture supernatants were collected at 24 hpi to estimate the viral copy number by qRT-PCR. The bar diagram indicates the $log_{10}$ of JEV RNA copy number/mL of the supernatants from different treatment conditions. (G) HEK293T cells were transfected with 1 µg of β-catenin plasmid for 24 h and then infected with JEV (MOI 0.1). Cells and culture supernatants were harvested at 24 hpi and subjected for downstream experiments. Western blot images representing the band intensities of the JEV-NS3 and total β-catenin proteins in empty vector and overexpression samples. (H, I) Bar Graphs representing the comparative expression levels of JEV-NS3 and total β-catenin between empty vector and overexpressed samples. GAPDH was used as an internal control. (J) Graph showing viral RNA as copy no./mL in β-catenin overexpressed HEK293T cells as relative to that of empty vector. Data are represented as mean ± SD and statistical significance was defined as P values $*\leq 0.03$, $**\leq 0.002$, $***\leq 0.0002$, $****<0.0001$ (n=3).

**Table S1: Details of the siRNA and primers sequence.**

| S.No. | Gene Name | Sequence |
|---|---|---|
| 1. | siRNA- β Catenin (Sense strand) | GUUGCUUGUUCGUGCACAUdTdT,<br>GCUUAUGGCAACCAAGAAAdTdT,<br>CAUGCAGUUGUAAACUUGAdTdT) |
| 2. | JEV-Envelope (E) | FP:5’-TTGACAATCATGGCAAACGA-3’<br>RP:5’-CCCAACTTGCGCTGAATAAT-3’ |
| 3. | GAPDH | FP:5’-GAGTCAACGGATTTGGTCGT-3’<br>RP:5’-GACAAGCTTCCCGTTCTCAG-3 |
| 4. | Cyclin D-1 | FP:5’- GGATGCTGGAGGTCTGCGAGGAAC-3'<br>RP:5’- GAGAGGAAGCGTGTGAGGCGGTAG-3' |

**Table S2: The percent possibility of complex formation between total β-catenin and JEV proteins.**

| | **Possibility of complex formation (%)** | | | |
|---|---|---|---|---|
| **Interaction with total β-catenin** | **Transient complex** | **permanent complex** | **Crystal Artifact** | **Free Energy Kcal/mol** |
| E | 92 | 1 | 7 | -12.8 |
| NS1 | 76 | 23 | 1 | -17.3 |
| NS3 helicase | 70 | 26 | 4 | -13.6 |
| NS5MTase | 32 | 65 | 3 | -11.4 |
| NS5RdRp | 92 | 7 | 0 | -16.7 |
